# High throughput detection of variation in single-cell whole transcriptome through streamlined scFAST-seq

**DOI:** 10.1101/2023.03.19.533382

**Authors:** Guoqin Sang, Jiaxin Chen, Meng Zhao, Huanhuan Shi, Jinhuan Han, Jiacheng Sun, Ying Guan, Xingyong Ma, Guangxin Zhang, Yuyan Gong, Yi Zhao, Shaozhuo Jiao

**Author notes:** Authors contributed equally to work (Guoqin Sang, Jiaxin Chen, Meng Zhao, Huanhuan Shi).

## Abstract

High-throughput single-cell full-length RNA sequencing is a powerful tool to explore the entire transcriptome, including non-polyadenylated transcripts. We developed a <u>s</u>ingle <u>c</u>ell <u>F</u>ull-length RN<u>A S</u>equence <u>T</u>ranscriptome <u>seq</u>uencing method (scFAST-seq), which combines semi-random primers with high reverse transcription efficiency, template switching and convenient rRNA removal methods, allowing the construction of full-length RNA libraries of up to 12,000 cells within 8 hours. Compared to regular 3’ scRNA-seq, scFAST-seq has similar sensitivity to mRNA detection, sequencing cost and experimental workflow. Moreover, scFAST-seq has clear advantages in detecting non-polyadenylated transcripts, covering longer transcript length, and identifying more splice junctions. In addition, scFAST-seq can more accurately predict the direction of cell differentiation by calculating RNA velocity. Furthermore, we demonstrated that scFAST-seq combined with target region enrichment can simultaneously identify somatic mutations and cellular status of individual tumor cells, which is valuable information for precision medicine.

## Background

Single-cell RNA sequencing (scRNA-seq) technology can reveal new insights into cell heterogeneity by analyzing transcriptomes in tens of thousands of individual cells. This makes it an important and powerful tool for understanding the mechanisms of life. However, current scRNA-seq methods primarily utilize oligo-dT primers to initiate reverse transcription from the polyadenylated tail of mRNAs [1], resulting in low sensitivity for detecting non-polyadenylated transcripts such as long non-coding RNAs (lncRNAs), histone mRNAs [2], ribosomal RNAs, circular RNAs (circRNAs), and enhancer RNAs (eRNAs) [3]. Moreover, high-throughput scRNA-seq methods have an obvious 3 ′- or 5 ′-end bias due to their short library fragments (∼300-700 base pairs), which must contain cell barcodes adjacent to the poly(A) tail or 5′ end of the transcripts. This bias limits their potential applications in detecting mutations or alternative splicing events that could occur at any position along the RNA strand.

However, several high-throughput scRNA-seq methods have been developed to overcome these limitations. By using long-read sequencing to analyze barcoded cDNA products derived from high-throughput scRNA-seq platforms, full-length RNA sequences can be detected to analyze alternative splicing events [4-7]. Nevertheless, the cost of long-read sequencing and the inability to detect non-polyadenylated transcripts hinder their widespread application. VASA-seq (Vast Transcriptome Analysis of Single Cells by dA-Tailing Technology) can capture both non-polyadenylated and polyadenylated transcripts and sequence them on low-cost short-read next-generation sequencing platforms without significant 3′ or 5′-end bias [8]. However, VASA-seq requires repetitive droplet generation and picoinjection, making it too complicated for most biological laboratories.

To overcome these challenges, we developed a <u>F</u>ull-length RN<u>A S</u>equence <u>T</u>ranscriptome sequencing method (scFAST-seq), which combines semi-random primers with high reverse transcription efficiency and convenient rRNA removal technologies during the cDNA amplification step. scFAST-seq has comparative sensitivity for transcript detection, sequencing cost and streamlined experiment processes to regular 3’ scRNA-seq, allowing for the construction of full-length RNA libraries with up to 12,000 cells within 8 hours. Our results suggest that the scFAST-seq has several advantages in cell clustering, RNA velocity inferring and non-coding transcripts identification. Notably, when combined with targeted region enrichment technology, the scFAST-seq can identify somatic mutations and cellular status of individual tumor cells, which is valuable information for precision medicine.

## Result

### The principle of scFAST-seq

To develop a widely accepted single-cell RNA sequencing (scRNA-seq) method that captures full-length transcripts, including non-polyadenylated transcripts, we considered using random primers instead of oligo-dT to initiate reverse transcription. This allows us to capture transcripts independent of their polyadenylated tails. We also chose to use water-in-oil droplets as a partitioning reaction assay to label transcripts from individual cells with unique cell barcodes and prevent cell-to-cell contamination. This approach can be conveniently adopted on similar platforms such as inDrop and 10X Genomics. Additionally, we prioritized using short-read next-generation sequencing platforms due to their low cost. Lastly, our method should have detection sensitivity comparable to its 3’ scRNA-seq counterparts.

However, when we replaced cell-barcoded oligo-dT primers with randomer-dN6 in our previously developed 3’ scRNA-seq method, we found that the detection sensitivity was reduced by more than three times compared to 3’ scRNA-seq (737 vs 271 median genes per cell at a comparable sequencing depth; see Supplemental Figure 1). Additionally, we observed substantial reads mapping to ribosomal and mitochondrial RNA, which increased sequencing costs. We determined that the reduced sensitivity was mainly due to the low reverse transcription efficiency of randomer-dN6. To improve reverse transcription initiation, we evaluated completely-random primers (dN9 and dN15) [9] and semi-random primers (5N3G/5N3T [10] or 12N7K (random 12N followed by 7bp with known sequence)and found that 12N7K provided the best sensitivity for transcript detection (Figure 1a and supplemental fig2).

**Figure 1.**
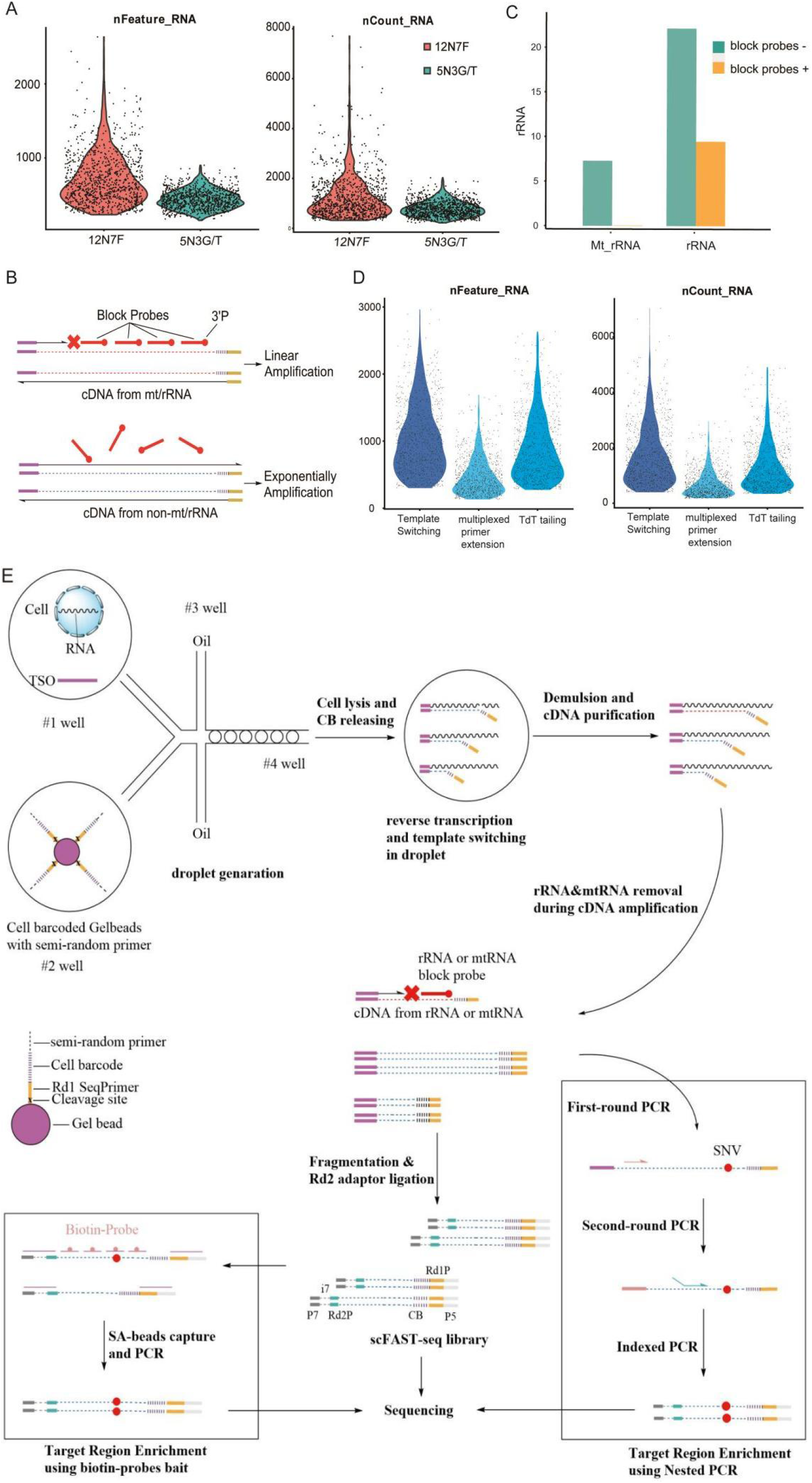
The principle of the scFAST-seq workflow. A. The comparison of detection sensitivity for number of genes and UMIs between 12N7K and 5N3G/T primers. B. Schematic illustration of the principle of ribosome and mtRNA removal. C. Comparison of ratio of sequencing reads mapped to ribosome or mitochondrial RNA in scFAST-seq data between libraries from cDNA amplified with or without block probes. D. Comparison of sensitivity to detect RNA among template switching, multiplexed primer extension and TdT mediated tailing of cDNA ends methods. E. scFAST-seq workflow schematic diagram with two target region enrichment methods.

Several technologies exist for depleting ribosomal RNA (rRNA) [11], including separating rRNAs using hybrid-specific antibodies [12] or magnetic streptavidin-coated beads[13], selectively degrading rRNA using RNase H[14] and treating with duplex-specific nuclease (DSN)[10, 15]. However, none of these methods can be seamlessly applied to droplet-based scRNA-seq methods.. Here We developed a convenient rRNA depletion method derived from PNA-mediated PCR clamping[16, 17]. As shown in Figure 1b, we designed probes with a 3’ non-extension blocker (3’ phosphorylation) that could perfectly hybridize with cDNA from rRNA/mitochondrial RNA (mtRNA) and mixed these probes into the cDNA amplification assay. During the annealing and elongation steps of PCR, the probes rapidly bind to cDNA derived from rRNA/mtRNA and inhibit strand elongation when using a polymerase without 5’→3’ exonuclease activity. Meanwhile, RNAs not hybridized with probes can be exponentially amplified. This difference in amplification efficiency leads to minimal rRNA percentage in the final library after several PCR cycles. As shown in Figure 1c, the proportion of ribosomal reads in total sequencing reads was effectively reduced from 30% to 10%, and the proportion of mtRNA reads was reduced from over 6% to less than 1%.

Given that semi-randomers can anneal and initiate reverse transcription at multiple sites on transcripts, multiple cDNA chains can be transcribed from a single RNA molecule. As a result, other methods for adding adaptors to the 3’ end of cDNA may produce higher sensitivity for RNA detection than template switching, which preferentially adds TSO sequences to the 3’ end of cDNA where the complementary RNA has a 5’ cap structure. The adaptor added to the 3’ end of cDNA serves as a PCR primer to exponentially amplify cDNA and generate enough product for library construction. We compared two methods, multiplexed primer extension [18, 19] and TdT-mediated tailing of cDNA ends [20], with template switching. Surprisingly, template switching had better performance in terms of cDNA yield and median genes per cell compared to the other two methods (Figure1D).

In conclusion, we developed a <u>F</u>ull-length RN<u>A S</u>equence <u>T</u>ranscriptome sequencing method (scFAST-seq) that can capture RNA independent of the polyadenylated tail of transcripts using a simple process (Figure1E). First, fresh cells were suspended in reverse transcription master mix and then encapsulated into droplets with cell-barcoded gel beads. Each gel bead was coupled with cleavable oligos consisting of a read1 seqprimer and a specific 17bp cell barcode in front of the 12N7K sequence. RNA released from each cell was reverse transcribed to cDNA and barcoded by the corresponding semi-randomers and tailed by template switching oligos within droplets. After breaking the droplets, pooled cDNA from all droplets was purified and amplified by PCR while amplification of cDNA derived from rRNA and mtRNA was inhibited by blocking probes. The PCR product was then fragmented and ligated with a read2 seqprimer before constructing the library by selectively amplifying DNA with cell barcodes using a unique double-indexed primer. Furthermore, we also developed targeted region sequencing methods based on scFAST-seq using nested PCR or biotin-probe bait enrichment to obtain high depth at low sequencing cost.

### Consistency and differences between scFAST-seq and 3’ scRNA-seq

To evaluate the technical performance of the scFAST-seq method, we performed scFAST-seq and 3’ scRNA-seq on mixtures of K562, A549, and HCC827 cell lines as well as breast cancer (BRCA), glioblastoma (GBM), mouse pancreatic cancer model (PAAD), and PBMC samples. As expected, scFAST-seq effectively suppressed the proportion of reads aligned to rRNA and mtRNA in total sequencing data, which was even lower than that of 3’ scRNA-seq (Figure 2A). The gene body coverage map further showed that compared to 3’ scRNA-seq with its obvious 3’ end bias, scFAST-seq had homogeneous coverage along the body of protein-coding genes (Figure 2B). Detection sensitivity was comparable between the two methods and the number of genes detected by scFAST-seq technology was even higher in cell line mixtures and PBMC samples (Figure 2C). By calculating the correlation of average gene expression in each specific cell type, we evaluated gene expression correlation in cancer samples and found that scFAST-seq showed a significant positive correlation with 3’ scRNA-seq techniques (P value <2.2e-16; cor >0.8; see Figure 2D). We then applied typical correlation analysis methods from the Seurat package to integrate and visualize both scFAST-seq and 3’ scRNA-seq data. We found that clustering and relative positions of cells in both types of data were highly overlapped in two-dimensional UMAPs (Figure 2E). Furthermore, analysis of fibroblast subtype proportions in pancreatic cancer and epithelial cell proportions in breast cancer showed that proportions of different subtypes were consistent between scFAST-seq and 3’ scRNA-seq for subtypes such as Luminal HS, Luminal AV, and Myoepithelial (Figure 2F). Finally, we analyzed copy number variation in single cells from both scFAST-seq and 3’ scRNA-seq using the inferCNV pipeline. In human glioma, breast cancer, and mouse pancreatic cancer samples we found little difference between copy number changes detected by both methods.

**Figure 2.**
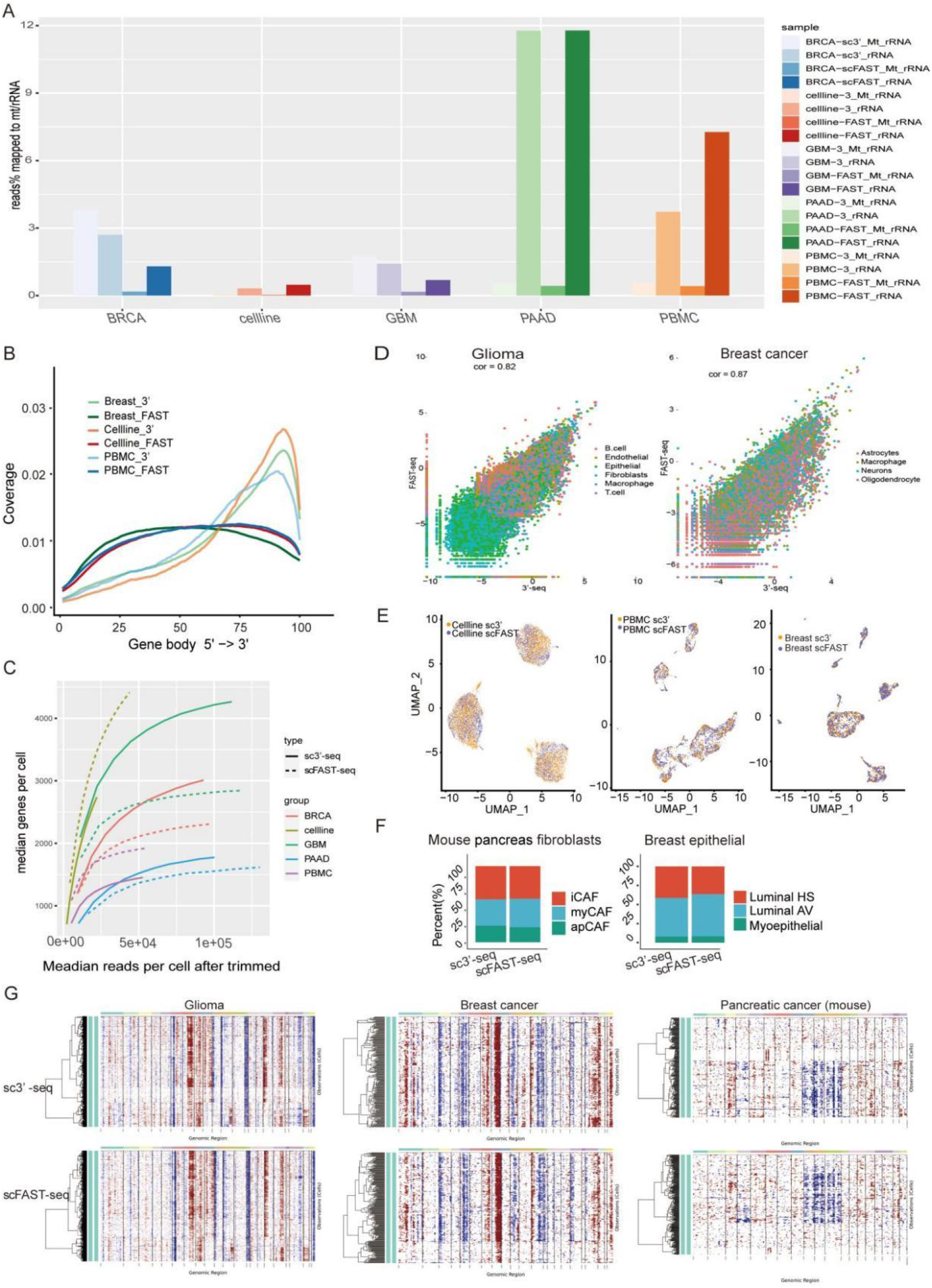
The consistency and differences between scFAST-seq and 3’-seq. A. Comparison of ratio of sequencing reads mapped to ribosome and mitochondrial RNA between 3’ scRNA-seq and scFAST-seq data in different tissues. B. Gene body coverage comparison. scFAST-seq showed even coverage, whereas 3’ scRNA-seq had a bias toward transcript termini in tissue, cell line mixture and PBMC samples. C. Plot showing median genes per cell as a function of downsampled sequencing depth in mean reads per cell between scFAST-seq and 3’ scRNA-seq data from breast cancer, cell lines mixture, gliblastoma, pancreatic cancer and PBMC. D. The gene expression correlation plot between FAST-seq and 3’-seq. A point presents a gene in specific cell type of glioma and breast cancer tissue. E. The overall consistency of cell clusters. The purple points are FAST-seq cells and the orange points are 3’-seq cells. F. The proportion of sub cell of mouse pancreas fibroblasts and breast epithelial. G. The copy number variation detected by inferCNV. The heatmap shows the relative intensity of gene expression on each chromosome, making it obvious which regions of the tumor genome are overexpressed or underexpressed.

### Advantages of scFAST-seq in transcript analysis

The scFAST-seq technology is superior to traditional 3’ scRNA-seq in terms of uniform coverage of gene body regions without terminal preference. This provides us with the opportunity to further study RNA characteristics and functions. It is well known that a single gene can produce multiple transcripts through splicing and that different transcripts can perform different biological functions within cells. We compared traditional 3’ scRNA-seq with scFAST-seq in terms of transcription characteristics and splice junctions. Firstly, we found that the ratio of transcription reads for lncRNA in GBM and BRCA samples using scFAST-seq was higher than with 3’ scRNA-seq. Additionally, the ratio of lncRNA transcription reads detected by scFAST-seq in PBMC was higher than with 3’ scRNA-seq (Figure 3A). To further demonstrate the higher capability to detect lncRNA, we compared published 3’ scRNA-seq data of lung cancer from 10X Genomics [21] and GEXSCOPE [22] platforms with scFAST-seq data of lung cancer (unpublished). We found that scFAST-seq could significantly detect more lncRNAs in lung cancer while maintaining equivalent expression levels for housekeeping genes (see Figure 3B). This result is not surprising because the 10X genomics or GEXSCOPE platform use oligo-dT primers either coupled to magnetic beads or hydrogel beads to capture poly(A)^+^ transcripts while non-poly(A) transcripts including lncRNAs were omited during reverse transcription.

**Figure 3.**
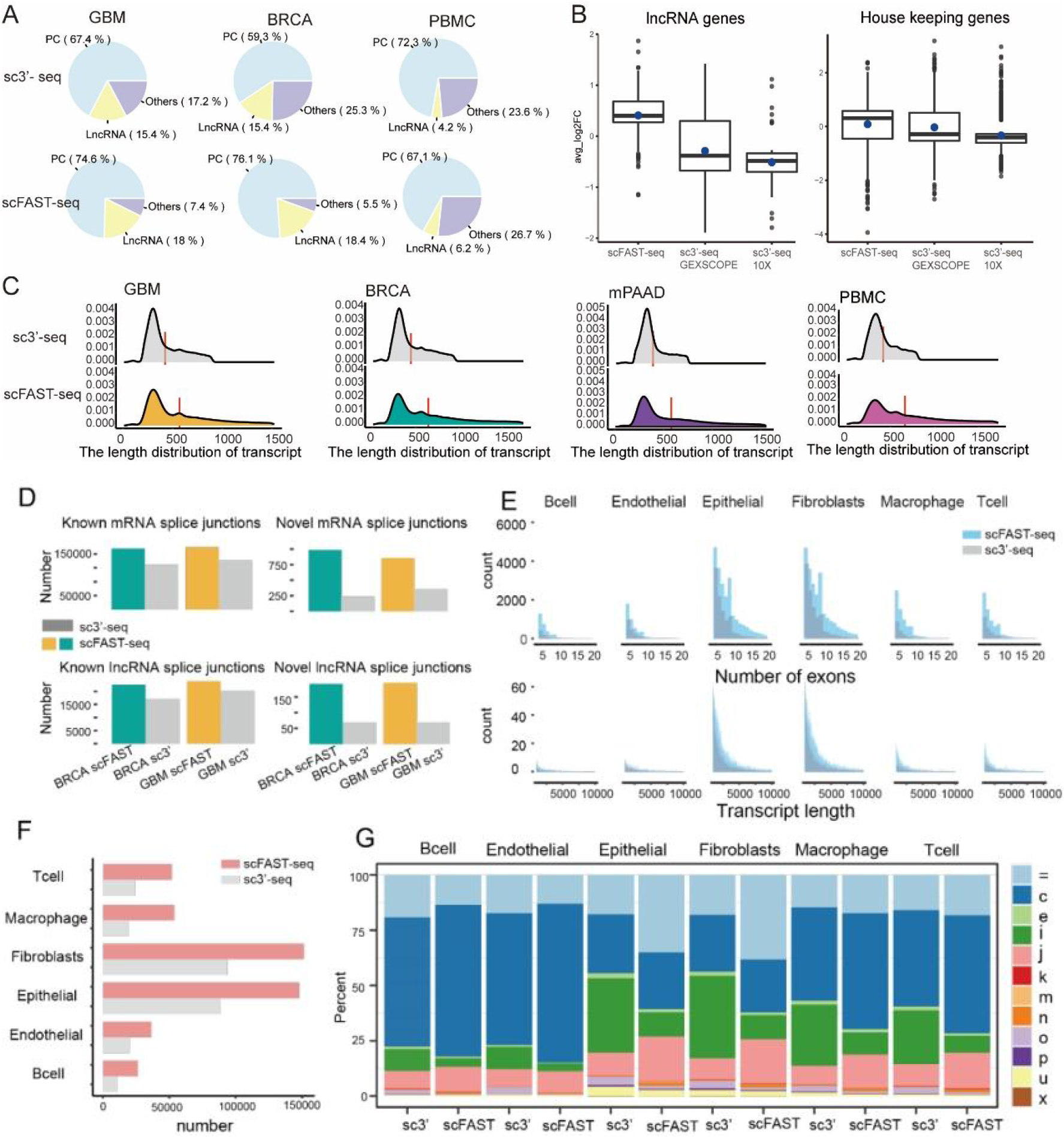
Advantage of transcript levels in scFAST-seq. A. The percentage of the number of mRNA and lncRNA transcript reads by 3’ sc-RNAseq or scFAST-seq. B. Box plot of expression levels of lncRNA and housekeeping genes between scRNA-seq and published 3’ scRNA-seq data from human lung cancer (log2 fold change calculated by FindMakrers function of Seurat software). We extract lncRNA genes from gencode gtf files, and used house keeping gene defined by RSeQC (https://sourceforge.net/projects/rseqc/files/BED/Human_Homo_sapiens/). To compare the expression levels of lncRNA and house keeping genes, we integrated our data with 3’ scRNA-seq data, and calculate the expression log fold change between the two data using Seurat software. C. The length distribution of the known gene transcripts. The red line location is median of length. D. A bar chart plot of the number of known and unknown splice junctions of recognized mRNAs and lncRNAs. E. Histogram of length and exon number distribution of reassembled transcripts. Blue represents the full length technique scFAST-seq and grey represents the sc3 ‘-seq. F. The number bar plot of splice junctions in different cell types of breast cancer. G. Proportion accumulation diagram of different types of transcripts. By assembling transcripts of each cell type separately, the different type of transcripts was annotated. For example, ‘=’ Complete match of intron chain; ‘c’ Contained; ‘j’ Potentially novel isoform (fragment) that shares at least one splice junction with a reference transcript.

We used the StringTie software to reassemble transcripts and obtain a new set of transcripts. Transcripts with an upper quartile length less than the length of all transcripts were selected for statistical analysis of their length distribution. We found that scFAST-seq could detect longer transcripts than 3’ scRNA-seq, which was consistent with the original assumptions and objectives of the scFAST-seq technique (Figure 3C). To find more sensitive new junctions, we used the STAR --twopassMode Basic method to enable more splice reads to map to new junctions. Statistically, we found that scFAST-seq discovered a greater number of known and new splice junctions (Figure 3D) than traditional 3’ scRNA-seq. We also performed this analysis at the cellular level and found that scFAST-seq detected more splice junctions (see Figure 3F) in each of six breast cancer cell types (see Figure 3F). Finally, by comparing reassembled transcripts with reference genomes, we annotated transcript types such as j, i, and u. Among six breast cancer cell types, scFAST-seq identified a significantly higher proportion of c and j transcripts than 3’ scRNA-seq with j transcripts defined as potential protein isoforms (see Figure 3G). In conclusion, scFAST-seq provided us with more opportunities and support to study variable splicing events and lncRNAs in disease at the single-cell level as well as more complex transcript types and functions.

### Accurately inferring the direction of T cell evolution by scFAST-seq

It has been shown that when mature mRNAs are expressed, a portion of immature transcripts are spliced. When gene expression increases, an instantaneous increase in the proportion of immature unspliced transcripts is observed within the cell. Conversely, when gene expression decreases, a higher proportion of spliced transcripts is seen for a short period of time. Therefore, we calculated the ratio of spliced to unspliced transcripts in our samples and found that scFAST-seq detected more unspliced transcripts in all three samples accounting for about twice as many as 3’ scRNA-seq (see Figure 4A).

**Figure 4.**
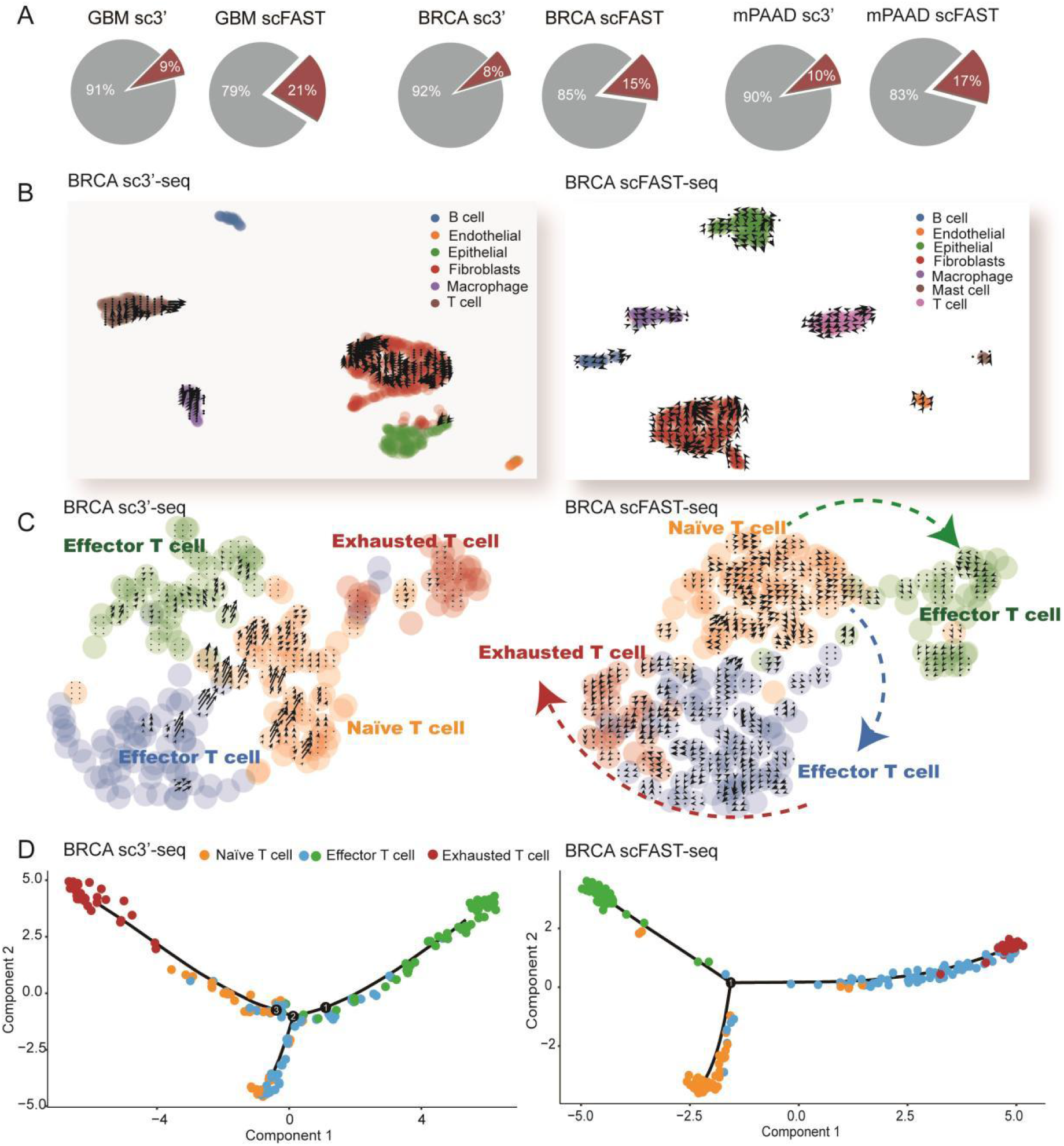
The evolution of T cell in breast cancer by scFAST-seq. A. The proportion of spliced and un-spliced RNA in scFAST-seq and 3’ sc-RNAseq. B. Cell cluster and RNA velocity of breast cancer sample in scFAST-seq and 3 ‘ sc-RNAseq. C. T cells can be divided into three states based on known cell markers, namely naive T cells, effector T cells, and exhausted T cells. The velocity of RNA was predicted by calculating the ratio of un-spliced to spliced RNA and the direction of T cell differentiation was deduced. D. The inference of differentiation trajectory by Monocle2. The sc3 ‘-seq is on the left and scFAST-seq on the right in Figure B-C.

In recent years, the concept of RNA velocity has been proposed in literature as an indicator to reveal dynamic changes in transcript abundance over time by evaluating the abundance of unspliced (nascent) and spliced (mature) mRNA. This is often used to study cell differentiation, lineage development and dynamic changes in tumor microenvironments. Based on our previous findings, we calculated the ratio of unspliced to spliced transcripts for each gene in single cells using both scFAST-seq and 3’ scRNA-seq data to reveal changes in gene expression and cell status in breast cancer samples. We used scVelo software to label cell state change directions on UMAPs according to RNA velocities calculated using ratios of unspliced-to-spliced counts for each gene. Our results showed that scFAST-seq could label directions for almost all cells while 3’ scRNA-seq could only label a small portion with epithelial endothelial and B cells remaining unlabeled (see Figure 4B)

In patients with chronic infections and cancer, T cells are continuously stimulated due to long-term exposure to persistent antigens and inflammation. This can lead to T cell exhaustion where exhausted T cells gradually lose their effector function and memory T cell features. In breast cancer samples, we separated T cells into three subtypes: ‘naïve’, ‘effector’ and ‘exhaustion and Treg’ according to previous studies. As shown in Figure 4C, the whole sequence technique can define the direction of differentiation for almost every T cell with predicted results conforming to known changes in the three states of T cells from initial to final exhaustion. However, 3’ scRNA-seq could only predict directions for a portion of cells with results not consistent with known directions of T cell differentiation (see Figure 4C). We also analyzed cell differentiation trajectories according to gene expression and found that scFAST-seq clearly described the branching trajectory of T cells from their origin to final exhaustion (see Figure 4D). These results indicate that scFAST-seq has obvious advantages in studying RNA velocity and inferring differentiation trajectories.

### Combining scFAST with target regions enrichment techniques to accurately detect gene mutations and fusion

Genome instability and mutations in driver genes are hallmarks of cancer and function-altering somatic mutations provide valuable information for basic cancer research and treatment [23]. Additionally, somatic mutations contribute to heterogeneity among tumor cells and alterations in tumorigenic signaling pathways. Therefore, there is an unmet need to detect somatic mutations in single cells. Given that sense somatic mutations can occur at any site within genes and must be translated into protein to gain function, scFAST-seq with its ability to detect full-length RNA is considered an ideal method for detecting mutations in single cells. To test this hypothesis, we equally mixed standard cell lines HCC827 (with EGFR 19del), A549 (with KRAS G12S) and K562 (with BCR-ABL fusion) and performed scFAST-seq to assess mutation detection sensitivity. As shown in Figure 5A, cell clusters were correctly identified as the three cell lines indicating minimal cross-contamination with scFAST-seq. However, only 6.77% of A549 cells had the KRAS G12S mutation and 30.53% of HCC827 cells had the EGFR 19del mutation (Figure 5A, left panel). Further analysis indicated that lower sequencing depth and coverage contributed to this low sensitivity of mutation detection. However, increasing transcriptome library sequencing data was not an option due to high costs so we developed two target region enrichment methods based on scFAST-seq to detect mutations with high sequencing depth at a lower cost.

**Figure 5.**
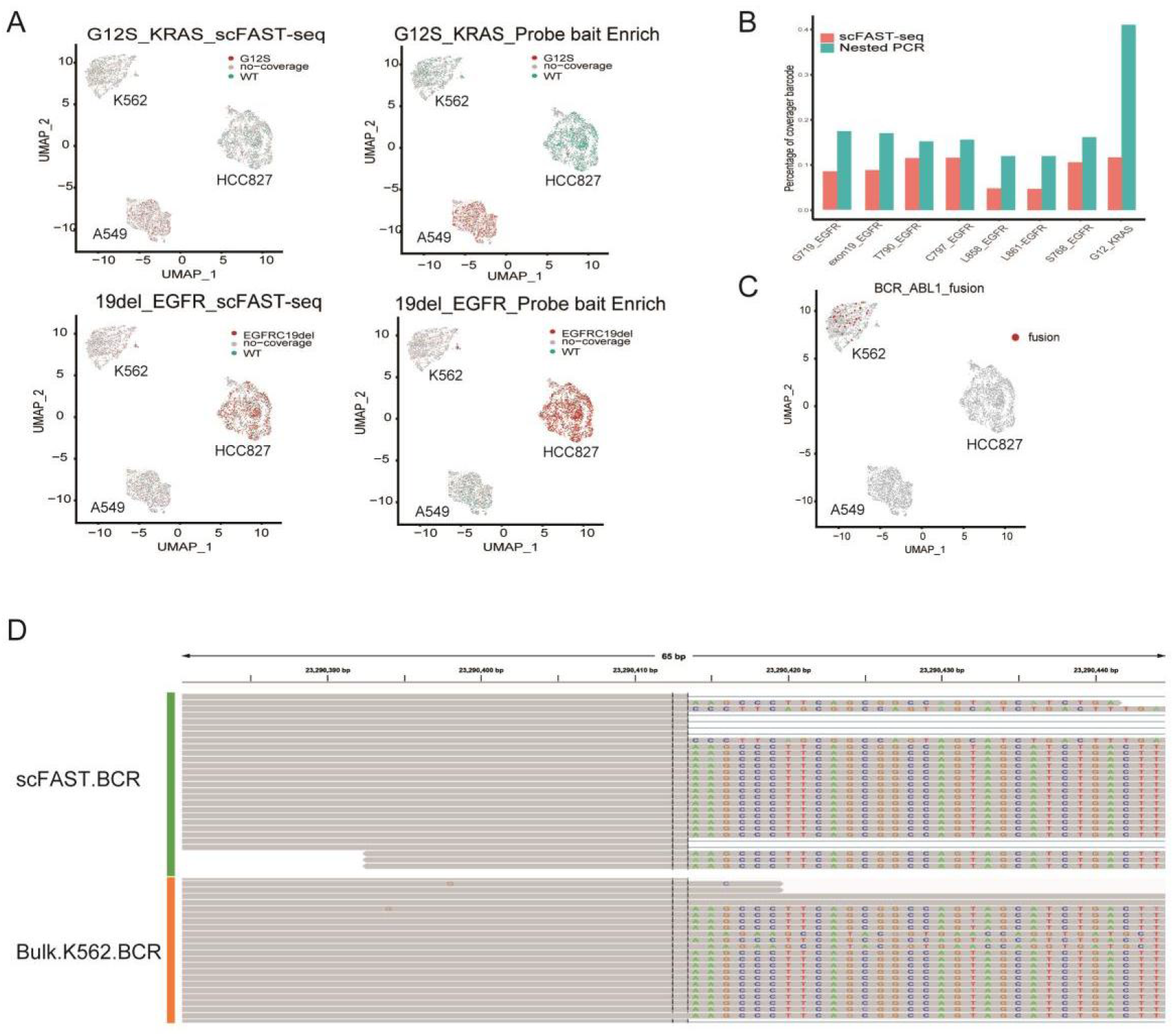
The mutation detection by scFAST-seq and panel enrichment. A. The clusters of subpopulations in A549, K562 and HCC827 cell mixture in scFAST-seq with (right panel) or without (left panel) biotin probes bait enrichment after running PCA and UMAP and the coverage of KRAS_G12S and EGFR_19del detected in each cluster. WT: wild type meaning there were reads covering this site but is non-mutation genotype. No coverage means no sequencing reads were detected covering this site. B. The different coverage rate of cells that had sequencing reads mapped to the corresponding sites of EGFR and KRAS genes between scFAST-seq with or without nested PCR enrichment. C. BCR-ABL fusion could be detected in K562 cluster by scFAST-seq with panel enrichment techniques. D. Visualization of the same breakpoint site of BCR in IGV between scFAST-seq and bulk sequencing.

Firstly, we assessed the biotin-probe bait enrichment method due to its high scalability from hundreds of genes to the entire human exome (Figure 1E, right panel) [24]. Compared with the low detection sensitivity of transcriptome-scale scFAST-seq, the G12S mutation could be detected in 28.50% of A549 cells using biotin-probe bait enrichment methods while the percentage for EGFR 19del increased from 30.53% to 70.01% in HCC827 cells (see Figure 5A). Next, we used nested PCR which has been applied in Cytoseq to detect low-abundance transcripts and rare cells (see Figure 1E, right panel) [25] to enrich fragments containing KRAS G12 site and EGFR hotspots from amplified cDNA. As shown in Figure 5B, fractions of cells with reads covering specific hotspots significantly increased by 2-3 times after nested PCR enrichment. We also detected some gene fusion sites using STAR-Fusion software based on genome alignment results with panel enrichment techniques. For example, we detected the BCR-ABL1 fusion gene in the K562 cell line (Figure 5C) although its detection rate was only 2.86%, significantly lower than G12S and 19del mutations. The breakpoint of BCR occurred at the same site as visualized in the Integrative Genomics Viewer between data from scFAST-seq with enrichment and bulk sequencing technology (Figure 5D).

Taken together, these results illustrate that combining scFAST-seq with target enrichment can significantly increase sensitivity for detecting mutations in exons at the single-cell level.

## Discussion

We developed scFAST-seq to provide a high-throughput and streamlined process for detecting tens of thousands of single-cell transcriptomes within 8 hours. By using barcoded semi-random primers to initiate reverse transcription, all exons independent of its location in RNA and non-polyadenylated transcripts have an equal opportunity to be analyzed. We noticed that 3 ′ scRNA-seq could also detect substantial amounts of lncRNA on droplet-based platforms. Further analysis revealed that oligo-dT primers could initiate reverse transcription at A-rich sequences located in the middle or at the 3′ tail of lncRNA. This enrichment of A-rich lncRNA can create detection bias in 3 ′ scRNA-seq, while the use of semi-random primers in scFAST-seq can alleviate this bias.

Despite its advantages, scFAST-seq still has some shortcomings that need to be optimized. For example, the detection rate of BCR-ABL fusion in K562 cells was lower than expected. Possible reasons for this include limited expression copies of fusion genes and lower efficiency of semi-random primers in capturing some regions of RNA without complementary sequences. We also noticed that the coverage rate of specific genes varies greatly depending on their expression levels. This is mainly because sequencing reads belonging to one UMI can assemble and cover a sequence of 500-1000bp. Thus, for genes with a length of 2000bp, at least 3 UMIs - but frequently 10 UMIs in practice - are needed to achieve 90% coverage of a specific gene in a single cell. This issue may be alleviated by increasing sensitivity through the recovery of more cDNA, as Seqwell-S^3 does[26].

In summary, scFAST-seq has several advantages over 3 ′ scRNA-seq. These include better detection of non-polyadenylated transcripts, longer coverage of transcripts, identification of more splice junctions, and more accurate prediction of cell differentiation direction. When combined with targeted region enrichment, scFAST-seq has greater potential to detect randomly occurring mutations in exons at the single-cell level. Overall, scFAST-seq is a desirable alternative to 3 ′ scRNA-seq - especially considering its comparable sensitivity for mRNA detection, sequencing cost and experimental workflow.

## Method

### Single cell suspension acquisition and counting

For preparation single cell suspensions, tissues were washed in RPMI-1640 medium (Hyclone) and cut into small pieces. MACS Tumour Dissociation Kit (Miltenyi Biotec) were used for enzymatic digestion for 30 min on a rotor at 37 °C according to the manufacturer ‘s instruction. After filtration with 40μm cell strainer and red cell removal, single cells were suspended in RPMI-1640 with 10% fetal bovine serum (GIBCO) and counted on automated counter(Countstar Rigel S2).

Cell line K562, A549 and HCC827 (purchased from ATCC) was cultured in a RPMI-1640 or DMEM complete medium (Hyclone) containing 10% fetal bovine serum (GIBCO) with 5% CO2 incubator at 37℃. After harvest from culture dishes, single cell suspensions were counted on automated counter(Countstar Rigel S2) and diluted into 1000 cells/μL for droplet generation.

#### Human Specimen

Surgical Tissues of breast cancer and glioma were obtained from volunteers who had undergone surgery at the Cancer Hospital, Chinese Academy of Medical Sciences between July and August 2022. PBMC were separated from blood of volunteers with 1.077g/ml ficoll. All volunteers have been well informed.

### Droplet Generation and reverse transcription

All the reagents and consumables used here were from commercial SeekOne^®^ DD single-cell 3’ Transcriptome Kit (SeekGene Biosciences) except gel beads. Unlike single cell 3’ gel beads with barcoded oligo-(dT)_30_ primer in kit, the semi-random primer of scFAST-seq was barcoded 12N7K with sequence CTACACGACGCTCTTCCGATCT(j)_17_(N)_12_TTGCTGT, where (j)_17_ represents the 17bp cell barcode sequences and (N)_12_ represents the random 12bp sequences (available as universal gel beads, SeekGene Biosciences). In some cases the primers coupled to gelbeads were 5N3G/T with sequences CT…CT-(j)_17_(N)_5_GGG and CT…CT-(j)_17_(N)_5_TTT, dN9 with sequence CT…CT-(j)_17_(N)_9_ or dN15 with sequence CT…CT-(j)_17_(N)_15_.

Typically 3K to 20K cells and water were added to the 46.8μL RT primix(20μL RT buffer, 18μL Indicator, 5.2μL RT Enzyme, 2μL TSO and 1.6μL Reducing buffer) for a total of 80µl. After pipette mixing 10 times, 78μL RT mixture were was loaded into #1 well. Gel beads with oligo-(dT)_30_ primer or random primer were loaded into #2 well followed with carrier oil into #3 well in chip. Then droplets were generated into #4 collection well within 5 minutes using SeekOne^®^ digital droplet instrument and transferred into a new PCR tube in where reverse transcription were performed by running the following procedure: Hot cover 85℃, 15 cycles (8℃ 12 s, 15℃ 45 s, 20 ℃ 45 s, 30 ℃ 30 s, 42℃ 3 minutes, temperature change rate 1.5℃/s); 85℃ for 5 minutes.

### cDNA amplification and rRNA/mtRNA removal

After emulsion breaking and purification, cDNA were added to the PCR assay containing 25μL 2×KAPA, 0.4μL cDNA Primer, and enzyme-free water with total 50μL. After 98℃ for 3 min, 13 cycles were performed (98℃ for 10 s, 63 ℃ for 15 s, 72 ℃ for 3 min) for 3’ scRNA-seq. To reduce the percentage of rRNA/mtRNA in the PCR products, 1.45μL block probes (final concentration 0.2 μ M each) were added to the assay and two-round PCR were performed following protocol: 98℃ 3 min, (98℃ 10 sec, 63℃ 15 sec, 72℃ 3 min) for 7cyles, 72℃ 5 min, 4℃ Hold. The product of first PCR round were purified with 0.6X SPRI beads and served as the template of the second PCR. The final product from the second round PCR was also purified with 0.6X DNA clean beads (Vazyme) and dissolved in 40μL Nuclease free water.

### TdT mediated tailing of cDNA ends

After emulsion breaking and purification, Tailing and ligation of T7 adaptor was performed to the 3’ end of cDNA according to manuscript of Scale ssDNA-seq lib Prep kit for illumina V2(10pg-200ng, RK20228, ABclonal). After second strand synthesis reaction and purification with 0.6X DNA clean beads, the products were amplified and purified with 0.6X DNA clean beads and used to fragment for library preparation as follows.

### Library preparation and sequencing

A quarter of the volume of the cDNA product was fragmented, end repaired and added with “A”. After size selection with 0.6x/0.2x DNA clean beads, the resulting product was ligated to the Illumina Truseq adapter using T4 DNA fast ligase. Ligation products were purified using 0.8x DNA clean beads and amplified in 50μL assay with 25μL 2x kapa HiFi HotStart Ready mix, 2μL 10μM index primers. After incubation at 98°C for 3 min, 13 PCR cycles were performed (98°C 20 s, 54°C 30 s, 72°C 20 s). The final product was size selected by 0.5x/0.3x DNA clean beads. The library was sequenced on Illumina Novaseq 6000 with PE150 strategy.

### Multiplexed primer extension

After emulsion breaking and purification, cDNA were added to the assay containing 5μL NEBuffer 2, 2μL 10mM dNTP, 1.5μL Klenow Fragment (3’ -> 5’ exo^-^, NEB), 5μL 100μM random primer (TCAGACGTGTGCTCTTCCGATCTNNNNNNNNN) for a total 50μL with a protocal: 25°C 10min, 37°C 20min, 50°C 10min. After purification with 1.8X DNA clean beads, the products was amplified for 10-13 cycles (98℃ 30 sec, 63℃ 1 min, 72℃ 1 min) with prime pairs (Forward: ACACTCTTTCCCTACACGACGCTCTTCCGATCT and Reverse: GTGACTGGAGTTCAGACGTGTGCTCTTCCGATCT). The amplified product was purified with 1X clean beads and amplified with indexed primer to obtain final libraries.

### Target Region Enrichment

For nested PCR, amplified cDNA were chosen to be the initial template for first PCR round using primers shown in supplemental tab1. After purification with 0.8X DNA clean beads, the products of second round PCR were used to perform indexed PCR to obtain the final enrichment libraries. For biotin-probes bait enrichment, the final libraries of scFAST-seq were hybridized with probes from Target Cap® Solid Tumor Fusion RNA-Cap Panel and captured by M270 streptavadin beads according to the manuscript (PD00696, Boke Bioscience). Followed by PCR using P5 and P7 primers, the final capture libraries were purified and analyzed on TapStation 4200 (Agilent).

### scFAST-seq and 3’ scRNA-seq data processing and cell cluster

Firstly, we identified and extracted the start location of barcode and UMI sequence from the paired end sequence data. And the length less than 50 of reads were discarded, the reads left behind were trimmed. We add the sequence of barcode and UMI to reads2 file for following genome mapping. Secondly, we used STAR[27] and Qulimap2[28]software for genome alignment and reads statistics while correcting UMI and filtering background. The original gene count and cell barcode matrix were obtained by featureCounts software[29]. And we used R package Seurat to filter low-quality genes and cells and normalized the filtered gene count matrix. Through Principal Component Analysis (PCA) method dimension reduction and Shared Nearest Neighbor (KNN) graph clustering, the cell clusters were obtained and we annotate the types according to the known cell markers from previous researches and public resources, such as Cellmarker database[30]. We used t-SNE and UMAP method for cell dimension reduction and visualization.

### Mutation detection

The R package inferCNV (https://github.com/broadinstitute/inferCNV) is used to infer copy number variation on our single cell RNA-Seq data by smoothed averages of gene windows model. We used STAR-Fusion software to detect fusion transcripts after the output of STAR alignment. The insertion and deletion (Indels) were called by a Java script VarScan2 behind the samtools mpileup process[31]. And we used ANNOVAR to annotate the functions of these mutations. Finally, the genome reads covering location regions are presented by IGV software.

### Expanding the reference transcriptome annotation

Firstly, we used the clean reads and mapped to the human reference genome (GRCh38) using STAR software with ‘twopassMode Basic’ parameter and the alignment results and splice junction information was obtained. Secondly, we used StringTie2[32] software to assemble transcripts with the aid of known reference annotation. The information of transcripts such as length, exon number, read counts and coverage score were calculated for further statistics and screening. The type of transcripts with “class code” was identified through comparing the reference by gffcompare[33] software, such as ‘=’ type represents the complete match of intron chain ; type ‘u’ represents unknown, intergenic transcripts; type ‘o’ represents generic exonic overlap with a reference transcript. The protentional novel lncRNA transcripts were selected according to some criterions[34]: (1) the class code of transcripts was “i, x, u”; (2) the length of transcripts was more than 200nt, the FPKM value was more than 0.5 and the coverage score was more than 1; (3) the transcripts were classified to lncRNA by CNCI[35] and CPC2[36] software that were used to predict protein-coding ability by sequence. Finally, the extra and novel transcripts were also included into initial transcriptome annotation file to other analysis. At the cell type level, we divided the clean reads according to the labeled cell types and got the cell type specific transcripts according to the above process.

### Determination of T cell differentiation trajectory in breast cancer

We used scVelo package to visualize the cell direction of evolution and RNA velocity[37].

The T cell in breast cancer sample was separated to three subtypes, including ‘naïve’, ‘effector’ and ‘exhaustion and Treg’. Every cell subtype highly expressed the marker genes reported in previous studies (‘naïve’: LEF1, CCR7; ‘effector’: GNLY, CX3CR1; ‘exhaustion and Treg’: LAYN, CXCL13, FOXP3, CTLA4)[38]. And then the barcode and BAM file were used to infer the RNA velocity that is estimated by unspliced and spliced transcripts in single-cell RNA sequencing and represents direction of cell change by velocity software and scVelo python package. And we also calculated the proportion of spliced and unspliced transcripts in BRCA, GBM and mPAAD samples. In addition, we used Monocle2[39] to depict the T cell trajectory by the expression of specific state genes.

## Supplemental Figures

**Supplemental Figure 1.**
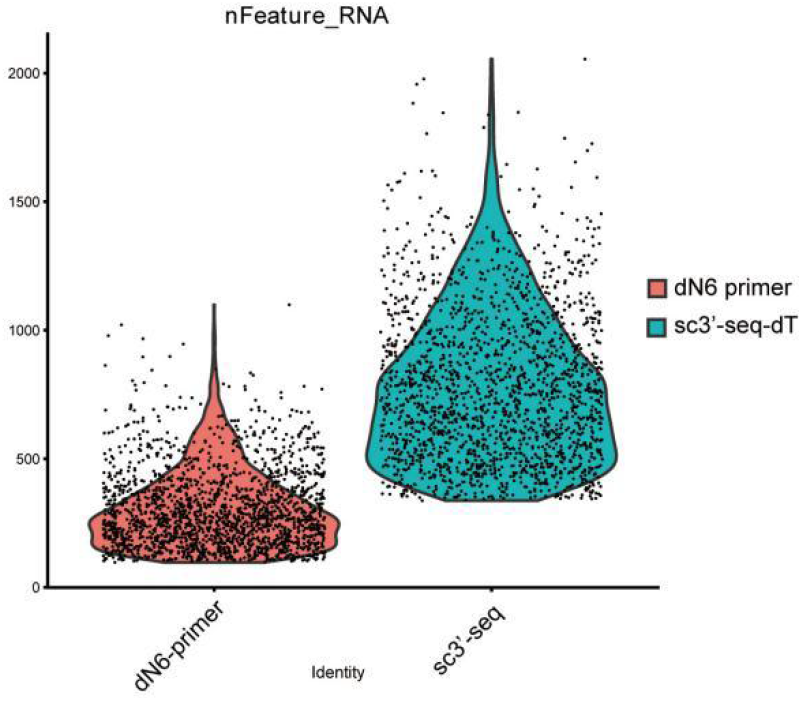

**Supplemental Figure 2.**
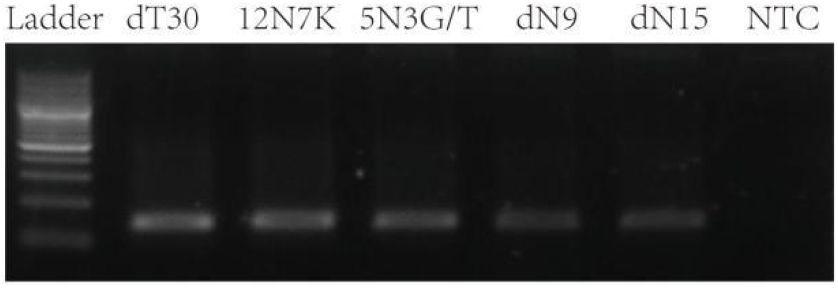

**Supplemental Table 1.**
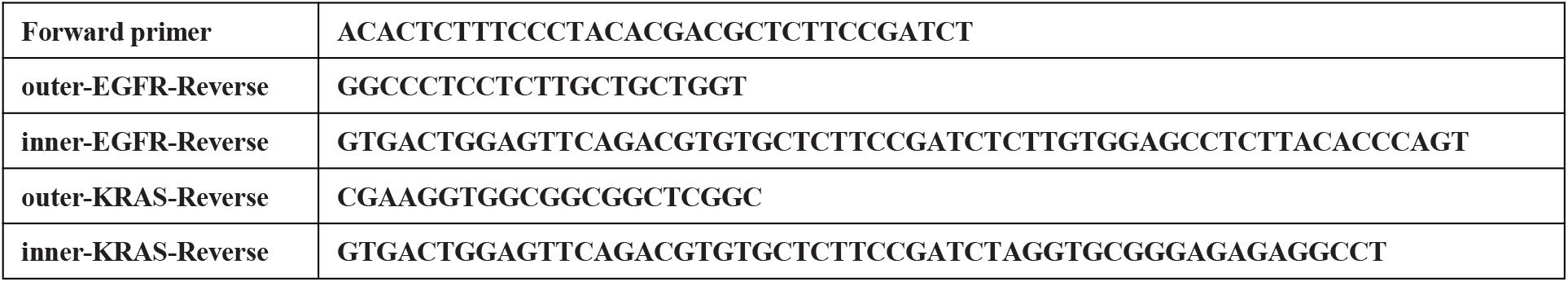

## Reference

1. Stuart, . and R. Sati a, ntegrative single-cell analysis. Nat Rev Genet, 2019. 20(5): p. 257–272.

2. Marzluff, W.F., E.J. Wagner, and R.J. Duronio, Metabolism and regulation of canonical histone mRNAs: life without a poly(A) tail. Nat Rev Genet, 2008. 9(11): p. 843–54.

3. Hayashi, ., et al., Single-cell full-length total RNA sequencing uncovers dynamics of recursive splicing and enhancer RNAs. Nat Commun, 2018. 9(1): p. 619.

4. upta, ., et al., Single-cell isoform RNA sequencing characterizes isoforms in thousands of cerebellar cells. Nat Biotechnol, 2018.

5. Singh, M., et al., High-throughput targeted long-read single cell sequencing reveals the clonal and transcriptional landscape of lymphocytes. Nat Commun, 2019. 10(1): p. 3120.

6. Lebrigand, K., et al., High throughput error corrected Nanopore single cell transcriptome sequencing. Nat Commun, 2020. 11(1): p. 4025.

7. Mamanova, L., et al., High-throughput full-length single-cell RNA-seq automation. Nat Protoc, 2021. 16(6): p. 2886–2915.

8. Salmen, F., et al., High-throughput total RNA sequencing in single cells using VASA-seq. Nat Biotechnol, 2022. 40(12): p. 1780–1793.

9. Stangegaard, M.,. H. Dufva, and M. Dufva, Reverse transcription using random pentadecamer primers increases yield and quality of resulting cDNA. Biotechniques, 2006. 40(5): p. 649–57.

10. Sheng, K., et al., Effective detection of variation in single-cell transcriptomes using MA Q-seq. Nat Methods, 2017. 14(3): p. 267–270.

11. Stark, R., M. Grzelak, and J. Hadfield, RNA sequencing: the teenage years. Nat Rev Genet, 2019. 20(11): p. 631–656.

12. Neil, D., H. Glowatz, and M. Schlumpberger, Ribosomal RNA depletion for efficient use of RNA-seq capacity. Curr Protoc Mol Biol, 2013. Chapter 4: p. Unit 4 19.

13. Culviner, P.H., C.K. Guegler, and M. Laub, A Simple, Cost-Effective, and Robust Method for rRNA Depletion in RNA-Sequencing Studies. mBio, 2020. 11(2).

14. Chen, Z. and X. Duan, Ribosomal RNA depletion for massively parallel bacterial RNA-sequencing applications. Methods Mol Biol, 2011. 733: p. 93–103.

15. Zhulidov, P.A., et al., Simple cDNA normalization using kamchatka crab duplex-specific nuclease. Nucleic Acids Res, 2004. 32(3): p. e37.

16. Cochet, ., et al., Selective PCR amplification of functional immunoglobulin light chain from hybridoma containing the aberrant M PC 21-derived V kappa by PNA-mediated PCR clamping. Biotechniques, 1999. 26(5): p. 818–20, 822.

17. von Wintzingerode, F., et al., Peptide nucleic acid-mediated PCR clamping as a useful supplement in the determination of microbial diversity. Applied and environmental microbiology, 2000. 66(2): p. 549–557.

18. Xu, H., et al., Detection of splice isoforms and rare intermediates using multiplexed primer extension sequencing. Nat Methods, 2019. 16(1): p. 55–58.

19. Shum, E., et al., Quantitation of mRNA Transcripts and Proteins Using the BD Rhapsody Single-Cell Analysis System. Adv Exp Med Biol, 2019. 1129: p. 63–79.

20. Schmidt, W.M. and M.W. Mueller, Controlled ribonucleotide tailing of cDNA ends (CR C) by terminal deoxynucleotidyl transferase: a new approach in PCR-mediated analysis of mRNA sequences. Nucleic Acids Res, 1996. 24(9): p. 1789–91.

21. Laughney, A.M., et al., Regenerative lineages and immune-mediated pruning in lung cancer metastasis. Nat Med, 2020. 26(2): p. 259–269.

22. Wu, F., et al., Single-cell profiling of tumor heterogeneity and the microenvironment in advanced non-small cell lung cancer. Nat Commun, 2021. 12(1): p. 2540.

23. Hanahan, D. and R.A. Weinberg, Hallmarks of cancer: the next generation. Cell, 2011. 144(5): p. 646–74.

24. Clark, M.J., et al., Performance comparison of exome DNA sequencing technologies. Nat Biotechnol, 2011. 29(10): p. 908–14.

25. Fan, H.C., K. Fu, and S.P. Fodor, Combinatorial labeling of single cells for gene expression cytometry. Science, 2015. 347(6222): p. 1258367.

26. Hughes, .K., et al., Highly Efficient, Massively-Parallel Single-Cell RNA-Seq Reveals Cellular States and Molecular Features of Human Skin Pathology. 2019.

27. Dobin, A., et al., S AR: ultrafast universal RNA-seq aligner. Bioinformatics, 2013. 29(1): p. 15–21.

28. Okonechnikov, K., A. Conesa, and F. Garcia-Alcalde, Qualimap 2: advanced multi-sample quality control for high-throughput sequencing data. Bioinformatics, 2016. 32(2): p. 292–4.

29. Liao, ., K. Smyth, and W. Shi, featureCounts: an efficient general purpose program for assigning sequence reads to genomic features. Bioinformatics, 2014. 30(7): p. 923–30.

30. Zhang, X., et al., CellMarker: a manually curated resource of cell markers in human and mouse. Nucleic Acids Res, 2019. 47(D1): p. D721–D728.

31. Koboldt, D.C., et al., VarScan 2: somatic mutation and copy number alteration discovery in cancer by exome sequencing. Genome Res, 2012. 22(3): p. 568–76.

32. Kovaka, S., et al., Transcriptome assembly from long-read RNA-seq alignments with String ie2. enome Biol, 2019. 20(1): p. 278.

33. Pertea, . and M. Pertea, FF Utilities: ffRead and ffCompare. F1000Res, 2020. 9.

34. Luo, H., et al., Single-cell Long Non-coding RNA Landscape of Cells in Human Cancer mmunity. enomics Proteomics Bioinformatics, 2021. 19(3): p. 377–393.

35. Sun, L., et al., Utilizing sequence intrinsic composition to classify protein-coding and long non-coding transcripts. Nucleic Acids Res, 2013. 41(17): p. e166.

36. Kang, .J., et al., CPC2: a fast and accurate coding potential calculator based on sequence intrinsic features. Nucleic Acids Res, 2017. 45(W1): p. W12–W16.

37. Bergen, V., et al., Generalizing RNA velocity to transient cell states through dynamical modeling. Nat Biotechnol, 2020. 38(12): p. 1408–1414.

38. Zheng, C., et al., Landscape of nfiltrating Cells in Liver Cancer Revealed by Single-Cell Sequencing. Cell, 2017. 169(7): p. 1342–1356 e16.

39. Qiu, X., et al., Reversed graph embedding resolves complex single-cell tra ectories. Nat Methods, 2017. 14(10): p. 979–982.

